# AlnC: An extensive database of long non-coding RNAs in Angiosperms

**DOI:** 10.1101/2021.02.04.429715

**Authors:** Ajeet Singh, AT Vivek, Shailesh Kumar

**Affiliations:** Bioinformatics Lab, National Institute of Plant Genome Research (NIPGR), Aruna Asaf Ali Marg, New Delhi 110067, India.

**Keywords:** Angiosperms, 1KP Project, lncRNAs, RNA-seq, sORFs

## Abstract

Long non-coding RNAs (lncRNAs) play a major role in diverse biological processes that are contemplated to have diverse regulatory roles in plants. While lncRNAs have been described and annotated in major plant species, many of plant species lack annotated lncRNAs despite the availability of transcriptome data in public repositories. In this study, we have identified and annotated the lncRNAs in the transcriptomic data from the 1000 plant genome project (1KP), and developed a user-friendly, open-access database called AlnC. This can be used for the exploration of lncRNAs in 682 Angiosperm plants. The current version of AlnC offers 10,855,598 annotated lncRNA transcripts across 809 sample tissues of diverse flowering plants. To enhance the AlnC interface, we have provided the features for browsing, searching and downloading of lncRNA data, interactive graphics, and online BLAST service. In addition to these information, each lncRNA record is annotated with possible Short open reading frames (e.g. sORFs) to facilitate the study of peptides encoded within lncRNAs. With this user-friendly interface, we assume that AlnC will serve as a rich source of lncRNAs that will contribute to small-and large-scale studies in a wide range of flowering plants.

## Introduction

Angiosperms (flowering plants) are land plants with more than 3,000,000 recorded species worldwide, comprising one of the most diverse group within the plant kingdom [1,2]. Most plants belonging to this group have been intensively studied in order to understand both the flowering and other major mechanisms. As a result, research in flowering plants has exploded with the advent of next generation sequencing leading to an improved picture of transcriptome, especially from the point of non-coding RNAs (ncRNAs). Of all recent studies in plant non-coding RNAs, long non-coding RNAs (lncRNAs) have been shown greater interest to study alongside miRNAs, which are typically more than 200 nucleotides with almost no protein coding capacity. There is also compelling evidence to lncRNAs functioning in multiple biochemical pathways of plants over the years [3–5]. Despite being in the spotlight, lncRNAs are yet to be annotated in a variety of plant species, despite the presence of data banks focused primarily on model plants and major crops. This research gap can be addressed by the use of RNA sequencing (RNA-seq) data as it provides enormous opportunities to understand transcriptome and classify potential lncRNAs [6,7]. Eventually, with the rise of transcriptome data in public repositories, thousands of plant lncRNAs were identified and maintained in a number of databases e.g. PLNlncRbase, CANTATAdb, GREENC, PLncDB v2.0, and lncRNAdb v2.0 [8–12]. The available lncRNA databases are limited to model plants and fewer other plants are another major concern, as information on the potential lncRNAs of many Angiosperms is still largely scarce and hinders the advancement of lncRNA research in several of these plants. Several plant species have been investigated for genome-wide lncRNAs using independent bioinformatics pipelines for annotation and archiving in databases, but several plants are still unexplored due to a complete lack of genome details. In order to address this problem in organisms with no genome data, various lncRNA methodologies are available, and are increasingly developed to improve lncRNA identification and annotation from *de novo* assembled transcripts [13,14].

In this current genomic era, it is essential to make lncRNA annotation models for diverse plant species in terms of improving our understanding of lncRNA biology. In parallel, the creation and maintenance of a stable lncRNA data repository is equally necessary if lncRNA biology is to be understood. [15,16]. In this research work, we have used large-scale transcriptome data to identify potential lncRNAs with three major goals. First, we anticipate to provide information on most potential lncRNAs of plant species with no available genome sequence. The second goal is to mediate the importance of unused large-scale datasets as an annotation source for thousands of lncRNAs. Lastly, to develop a database to act as a catalyst to promote lncRNA investigations in Angiosperms, and to provide a one-stop data access platform for lncRNA researchers. We therefore capitalised on large-scale transcriptomic data of 682 angiosperms from 1 Kp as this enabled us to annotate 10,855,598 lncRNAs. The results of the study have been organised and deposited in a user-friendly web-interface, the Angiosperm LncRNA Catalog (AlnC) with a plan to update periodically on the basis of new knowledge and an expansion in the number of species of Angiosperm in the future.

## Material and Methods

### Data collection and systematic lncRNA identification

We have downloaded the *de novo* assembled transcripts of 682 plant species belonging to Angiosperms from 1kp project (http://www.onekp.com/public_data.html), and identified the potential lncRNAs [17]. For each species, lncRNAs were identified from each library relying on a bioinformatics pipeline previously exploited by Singh et al., 2017, and transcripts longer than 200 bp were retained that are not overlapping with protein-coding gene models (Fig 1(A))[18]. First, we excluded potential coding transcripts (translated proteins of matched orthogroups derived from annotated plant genomes) from assembled transcripts set for that species [19]. Further, protein-coding transcripts were discarded by using PLncPRO (python prediction.py -p plncpro-result-file -i sequence.fa-m models/<monocot or dicot>.model -o plncpro-out -d lib/blastdb/swiss-protDB -t 15 - r) on the basis of a BLAST approach with the curated list of Swiss-Prot proteins [20]. In the final filtering step, high-confident lncRNA transcripts were extracted by setting a minimum length threshold of 200 nt, and a non-coding probability score of 0.8 (python predstoseq.py -f sequence.fa -o output-file -p plncpro-result-file -l 0 -s 0.8 --min 200).

**Fig 1.**
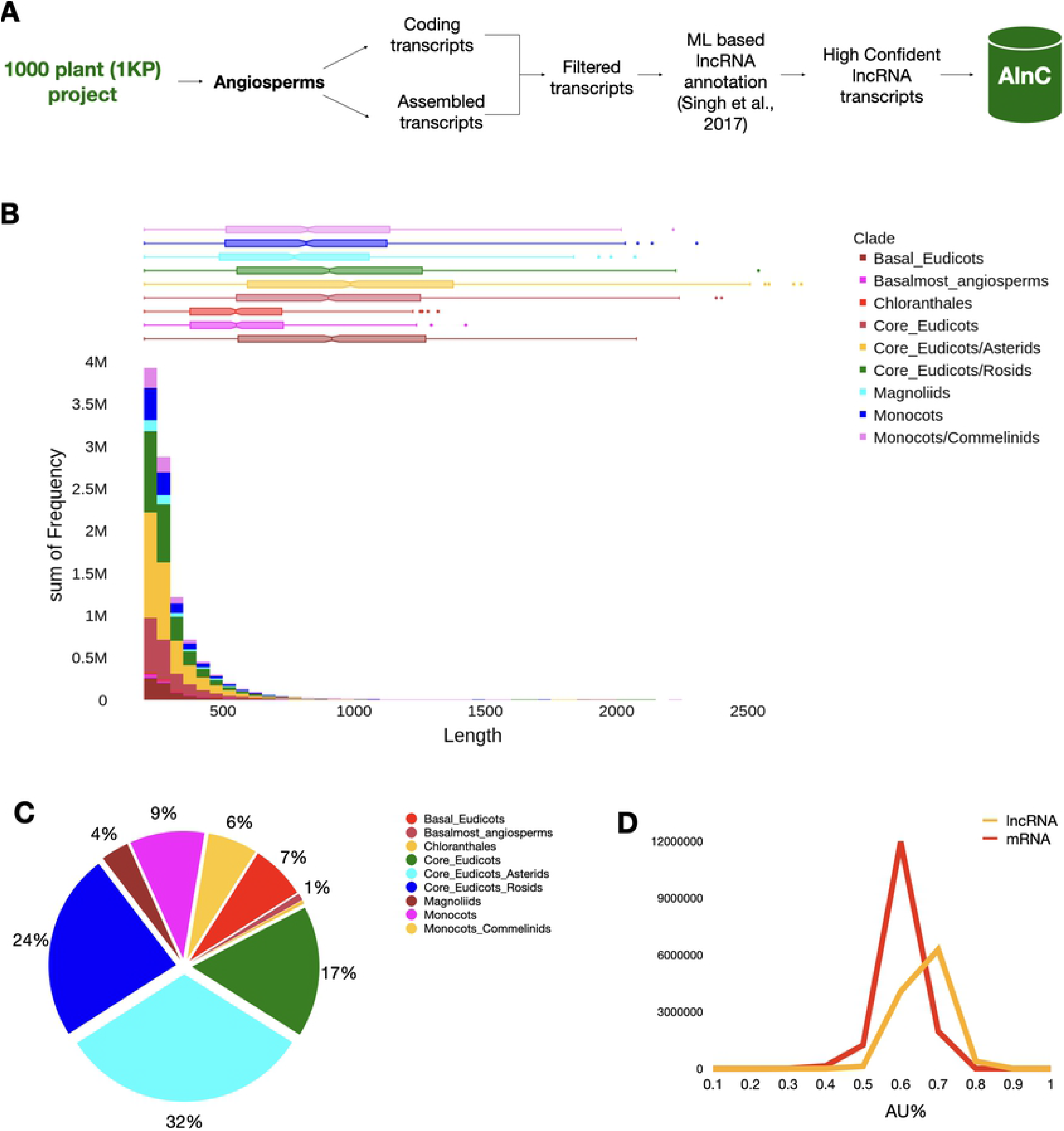
Overview of the lncRNAs in AlnC. (A) Systematic workflow adopted to annotate potential lncRNAs of all flowering plants available from the 1KP project, (B) Length-wise distribution of lncRNAs across clades, (C) Pie chart represents the percentage of lncRNA entries in AlnC, (D) Percentage composition of AU content in protein-coding transcripts and lncRNAs in AlnC.

Fig 1. (A) Systematic workflow adopted to annotate potential lncRNAs of all flowering plants available from the 1KP project.

### Database construction and implementation

After the compilation of all the information regarding lncRNAs, AlnC web interface was developed using Hypertext Mark-up Language (HTML), Cascading Style Sheets (CSS), Structured Query Language, Java scripting language, PERL and Hypertext Preprocessor on Apache Hypertext Transfer Protocol server. All lncRNAs available at AlnC, and other related annotations have been handled by a relational database set up with MySQL. A stand-alone BLAST (v2.11.0) tool is installed for online similarity search feature. ViennaRNA (v2.4.16) and ORFfinder (v0.4.3) tools are used for the secondary structure visualization, and identification of Small open reading frames (sORFs) respectively [21–23].

## Results and Discussion

### Data content in AlnC

We have identified a total of 10,855,598 lncRNAs from 809 RNA-seq samples of Angiosperms, available at 1kp project webpage (http://www.onekp.com/public_data.html). All the related numbers of total transcripts, filtered transcripts, lncRNAs, and filtered lncRNAs identified in each plant species are mentioned at the ‘Statistics’ section (http://nipgr.ac.in/AlnC/stat-data.php) of AlnC webpage. At AlnC, we have organised and compiled this catalogue of 10,855,598 lncRNAs from 809 samples available in the 1 KP project, which functions as a comprehensive catalogue of potential lncRNAs in 682 flowering plants. No other data repositories on lncRNAs on this scale exist, and most lncRNAs of the species included in AlnC belong to poorly studied taxa, rendering AlnC of wide interest among plant researchers (Fig 1(B & C)). At AlnC, the length of newly identified lncRNAs of flowering plants across clades have the size ranges from 200 to 7633 nt with an average length of 405 nt (Fig 1(C)). The median length of the identified lncRNAs is smaller than the median length of the coding sequences (Fig 1(B) and 3(E)). Moreover, most lncRNAs (66%) were less than 400 bp in length whereas only 2.9% lncRNAs were more than 1000 bp in length. Our attempts to identify ortholog relationships using BLASTN search reported large hits, however, we could only identify a moderate number of annotated lncRNAs at AlnC, identical to those stored in other databases reasoning to less conservation and apparent differences in the number of lncRNAs, but it also suggest certain significant sequence similarities across species. We have included the significant hits to NONCODE lncRNAs (384) and experimentally validated lncRNAs available in Plncdb (6) in Table S1 [11,24]. Further, the AU content of lncRNAs, available at AlnC, varied from 50-90% with an average of 75% in comparison to the coding transcripts which ranged 30-80% with an average of 60% (Fig 1(D)). Most lncRNAs contained more than 75% AU content, and the analysis implies the richness of AU than that in coding sequences [25–27]. A brief description of the lncRNA entries available at AlnC is represented in the Fig 1 and Table 1. The relationship analysis of annotated lncRNA with assembled transcripts and protein-coding transcripts yielded two distinct clusters of monocot and dicots, respectively (Fig 2). These clusters indicates the proportional discovery rate of lncRNAs with respect to the 1kp assembled transcripts, and also suggests the ratio of total transcripts to coding transcripts is smaller in dicot species whereas the reverse in case of monocots.

**Table 1:** Summary of annotated lncRNAs across higher-level clades in AlnC database.

| Clade | No. of Species | Number of Samples | Number of lncRNAs |
| --- | --- | --- | --- |
| Basal Eudicots | 33 | 55 | 835185 |
| Basalmost angiosperms | 8 | 8 | 281514 |
| Chloranthales | 2 | 2 | 46074 |
| Core Eudicots | 95 | 116 | 1828555 |
| Core Eudicots/Asterids | 217 | 242 | 3656812 |
| Core Eudicots/Rosids | 201 | 250 | 2627218 |
| Magnoliids | 26 | 27 | 383010 |
| Monocots | 61 | 64 | 829121 |
| Monocots/Commelinids | 39 | 45 | 568109 |
| Total | 682 | 809 | 10855598 |

**Fig 2.**
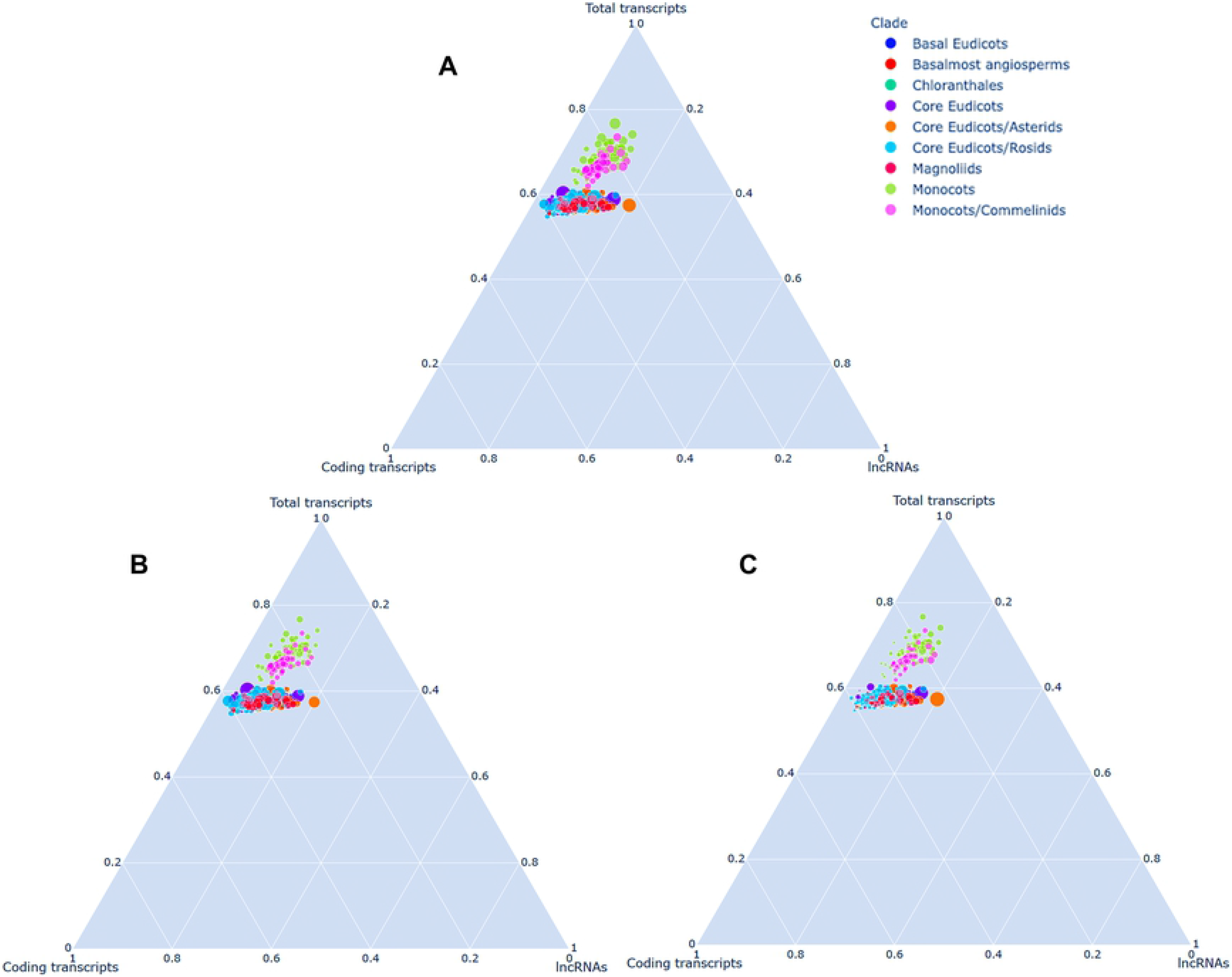
Ternary plot of AlnC lncRNAs, protein-coding and 1Kp assembled transcripts across clades. The bubble size represents the size of 1Kp assembled transcripts, protein-coding and AlnC lncRNAs in (A), (B) and (C), respectively.

Fig 1. (B) Length-wise distribution of lncRNAs across clades, (C) Pie chart represents the percentage of lncRNA entries in AlnC, (D) Percentage composition of AU content in protein-coding transcripts and lncRNAs in AlnC.

**Fig 2. Ternary plot of AlnC lncRNAs, protein-coding and 1Kp assembled transcripts across clades**. The bubble size represents the size of 1Kp assembled transcripts, protein-coding and AlnC lncRNAs in (A), (B) and (C), respectively.

### Modules available at AlnC

#### Search options

Current release of AlnC provides two search interfaces; (a) Simple Search, and (b) Advanced Search (Fig 3(A)). ‘Simple Search’ allows the user to quickly search the lncRNAs on the basis of taxonomic rank (Clade, Order, Family, Species), and non-coding probability score (min: 0.8; max: 1.0), while ‘Advanced Search’ provides enhanced query functionality using logical operators (AND/OR/>=/<=). Moreover, the ‘Advanced Search’ feature helps the users to search and apply filters to query a wide range of features and metadata fields e.g. Species, Tissue, lncRNA length, AlnC ID, and Non-coding probability. Consequently, a list of potential lncRNAs can be displayed at the result page according to the user defined queries, with the options to download and save the results. The search results shows the entries of the chosen species, covering the basic meta details of the 1kp sample code with links describing the sample information, including sample preparation, sample supplier, sample extractor, tissue type NCBI ID showing experimental sample library and run data, source transcript ID from which lncRNA is annotated, lncRNA length, and non-coding probability score (Fig 3(B)). The users can further explore the AlnC ID link to view and navigate at the detailed record of the lncRNA with primary lncRNA features, secondary structure, and other useful information e.g. ‘lncRNA Information’.

**Fig 3.**
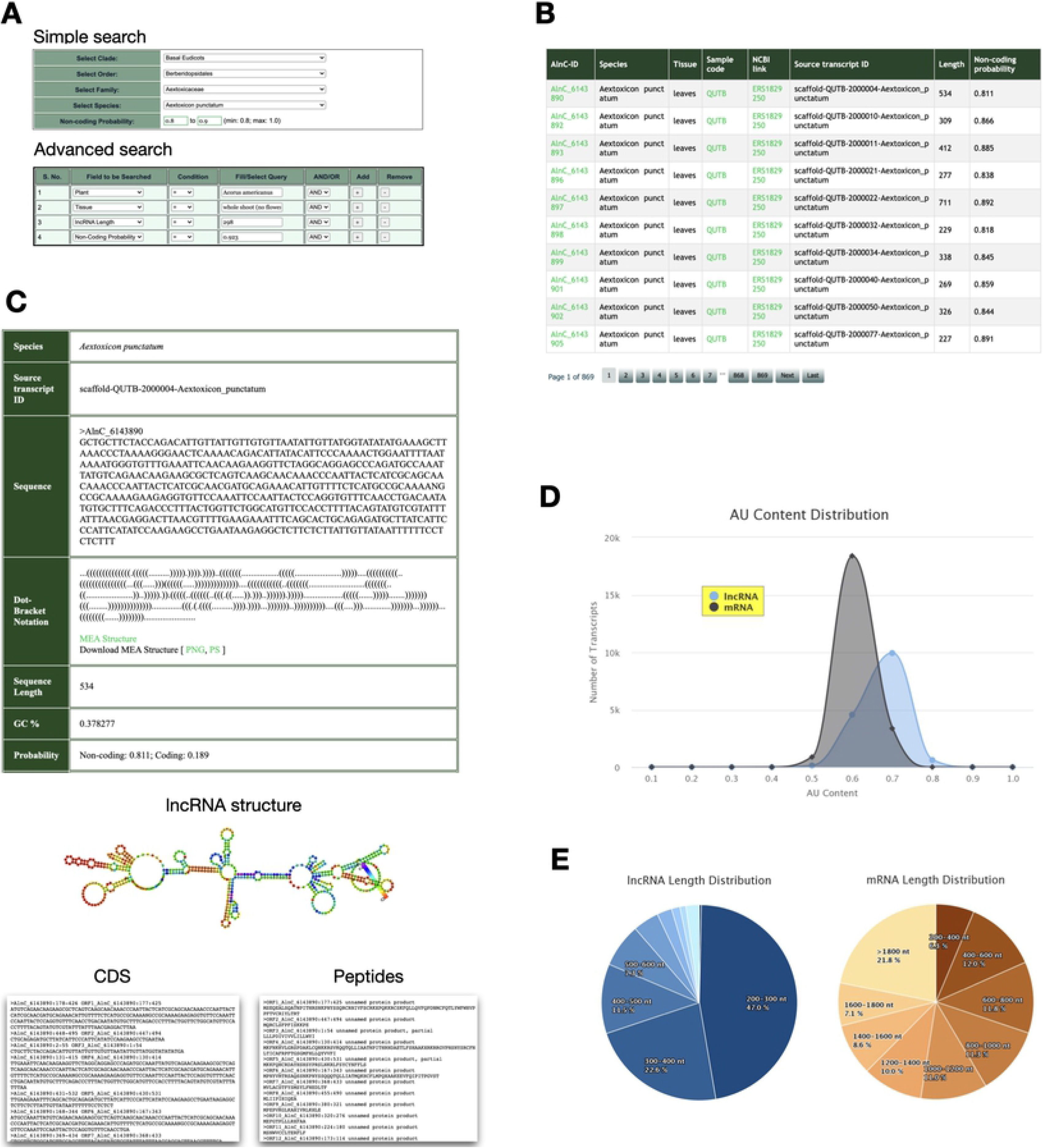
Screenshots of AlnC web interface. (A) Interface of simple search and advanced search modules, (B) Results page showing a table view of all lncRNA entries of the plant species *Aextoxicon punctatum,* (C) Sequence features including primary sequence information, secondary structure and possible sORFs as well peptides are displayed for lncRNA entry AlnC_6143890, (D) Percentage composition of AU content lncRNAs relative to mRNAs, (E) Pie chart representation of length-wise distribution of *Aextoxicon punctatum* lncRNAs and mRNAs.

Fig 3. (A) Interface of simple search and advanced search modules, (B) Results page showing a table view of all lncRNA entries of the plant species *Aextoxicon punctatum.*

Each ‘lncRNA Information’ page of AlnC enables the user to access to basic information of selected lncRNA with the information of structure (Fig 3(C)). This detail page is divided into two parts; the first section contains basic details of the lncRNA sequence (including the species name of the annotated lncRNA, the source transcript ID, length, GC content and coding/non-probability), and details of the secondary structure in dot-bracket notation with a provision for the user to view and download the structures in PNG or PS format. At the second section, user can also explore the ORFs and conceptual translation products of lncRNA sequence. All the associated annotations of the concerned lncRNA entry can be downloaded in a tabular form.

Fig 3. (C) Sequence features including primary sequence information, secondary structure and possible sORFs as well peptides are displayed for lncRNA entry AlnC_6143890.

#### BLAST module

BLAST feature is very helpful to find regions of similarity between the user provided input sequence and lncRNAs available at AlnC by using BLASTN with the option to change Expect value (E value). User can perform BLAST search against the database of a particular Clade, Order, Family, and Species. We have categorise the lncRNAs accordingly to avoid the memory intensive search against the full dataset available at AlnC. The BLAST search output includes pairwise alignment and a report with BLAST hits based upon alignment scores and other measures of statistical significance. After the BLAST run, user can directly go at the entry of any particular lncRNA showing similarly against the query sequence by a clink on the AlnC ID present in the BLAST results.

#### LncRNA vs mRNA module

This module of AlnC database allows the users to visualize and compare length-wise distribution, and AU% in annotated lncRNA transcripts and protein-coding transcripts in multiple samples of each species available at this database (Fig 3(D) and 3(E)). This page also enables the user to download the images of this comparative analysis in multiple file formats.

Fig 3. (D) Percentage composition of AU content lncRNAs relative to mRNAs, (E) Pie chart representation of length-wise distribution of *Aextoxicon punctatum* lncRNAs and mRNAs.

#### Download AlnC data

A this module, user can select the lncRNAs sequences for a particular Clade, Order, Family and Species, and download the complete sets from the AlnC archive. Complete AlnC collection can be found on the download module. Here users have access to both the hierarchical bulk download and the species-wise download in the FASTA file format.

### Submit Data

As there are sizeable researchers working on several flowering plants, we created a user form to submit any information or data with regard to lncRNAs of angiosperm species. This will allow us to develop, upgrade and manage AlnC on a regular basis. The related lncRNA data, research results and publications can be submitted using the form provided on the ‘Submit data’ page (http://nipgr.ac.in/AlnC/submit_data.php) of AlnC. If the submitted data found to be relevant, the received data will be curated manually and appended to AlnC. All submitted lncRNA data will be processed in compliance with the AlnC standard pipeline mentioned in Fig 1(A) and manually curated. We encourage submissions to AlnC curators as this will drive our plans to include additional species in the future.

### Conclusion and future prospects

In this research work, we have used the transcriptome data of 682 flowering plants, most of which had no genomic information and/or no documented lncRNA studies prior to this work. In our opinion, this is a key feature of AlnC, which was constructed with the primary objective of promoting the investigation of lncRNA in various Angiosperm species. The analysis workflow used in this study was optimized for plants and could be used for RNA-seq-based lncRNA identification for any other plant species as well. The workflow includes a ML based bioinformatics pipeline to identify high confident lncRNAs across distinct Angiosperm species, which significantly differs from other annotation approaches in lncRNA databases such CATATAdb [9]. Our approach yielded a large number of putative lncRNA transcripts, and all the annotated lncRNA transcripts were organised and catalogued in the AlnC web interface. It stores information of 10,855,598 lncRNA loci derived from 809 samples and provides a user-friendly platform for browsing, searching and accessing all annotated lncRNAs through simple and interactive web pages. AlnC includes lncRNAs with evidence of non-coding RNA probability score, and allows further exploration of sORFs alongside other primary lncRNA features, thus providing researchers with functional capability to leverage on AlnC data, and information on their individual projects using our web interface. With this research work, we have attempted to develop a first-ever database covering the largest number of confident lncRNA entries of wide-range plant species, including those with no information on lncRNA of any sort. Although, it is clear that several plants belonging to Angiosperms are still to be discovered and transcriptomes are waiting to be studied by individual research groups, AlnC will be continuously updated. At the same time, AlnC will strive to periodically search freely accessible databases, and other forms of documentation to collect useful information for annotated lncRNAs, and add additional functionality to enhance user engagement. It is our intention that AlnC will move forward to provide new databases representing additional species as well as to fine-tune, and optimise the currently available annotations. We will also aim to focus on the new pipeline to develop lncRNA annotations as the plant lncRNA biology research progresses. All in all, AlnC will strive to continue to be in line with the lncRNA community, and remain to serve useful lncRNA data in future.

## Acknowledgement

The authors are thankful to the Department of Biotechnology (DBT)-eLibrary Consortium, India, for providing access to e-resources. A.S and A.T.V are thankful to the Council of Scientific and Industrial Research and the DBT for research fellowships, respectively. Also, the authors acknowledge the Distributed Information Sub-Centers of DBT at the National Institute of Plant Genome Research (NIPGR).

## Author Contributions

A.S. and AT V. analysed the data and developed the web interface of this database. A.S., AT V., and SK wrote the manuscript. S.K. conceived the idea and coordinated the project. S.K. agrees to serve as the author responsible for contact and ensures communication.

## Supporting information

**Table S1:** List of significant hits of lncRNAs at AlnC, to lncRNAs available at NONCODE and PlncDB databases.

